# *Wolbachia* introgression in *Ae. aegypti* is accompanied by variable loss – a multi-country assessment

**DOI:** 10.1101/2024.08.21.608881

**Authors:** Kimberley R. Dainty, Etiene C. Pacidônio, Elvina Lee, Peter Kyrylos, Nathan Baran, Eleonora Kay, Yi Dong, Sofia B. Pinto, Gabriel S. Ribeiro, Alexander Uribe, Jovany Barajas, Scott L. O’Neill, Ivan D. Velez, Luciano A. Moreira, Cameron P. Simmons, Heather A. Flores

## Abstract

The *w*Mel and *w*AlbB strains of the bacterial endosymbiont *Wolbachia* are being introgressed into *Aedes aegypti* populations as a biocontrol method to reduce the transmission of medically important arboviruses. Successful introgression of *Wolbachia* relies on both persistence of *Wolbachia* throughout the host life cycle and a high fidelity of maternal transmission of *Wolbachia* between generations. *w*Mel has been introgressed into field populations in 14 countries to date. Monitoring of field sites has shown that *w*Mel prevalence can fluctuate substantially over time, prompting concerns this could lead to reduced efficacy of the biocontrol method. To explore the fidelity of *w*Mel persistence and transmission, we developed molecular methods to measure the prevalence of *Ae. aegypti* negative for *Wolbachia* infection but carrying the “founder” mitochondrial haplotype of the single female first transinfected. As all released *w*Mel-infected mosquitoes and any subsequent offspring will carry this founder mitochondrial haplotype, any mosquitoes with this mitochondrial haplotype and without *w*Mel indicate that *w*Mel was lost from this lineage at some point. We observed loss of *w*Mel ranging from 0 to 20.4% measured at various time intervals after *w*Mel-infected mosquito releases in five different countries. Despite some field sites showing *Wolbachia* loss, overall *Wolbachia* prevalence was sustained during the time periods studied. We then employed laboratory studies to explore factors that could contribute to the loss of *w*Mel. Surprisingly, near-perfect maternal transmission was measured across laboratory conditions of early blood feeding, starvation, and salinity. Collectively, these findings underscore that although *w*Mel transmission can be imperfect it does not necessarily undermine population-level establishment, providing encouragement that the intervention will be robust in most dengue-endemic environments.

## Introduction

The prevalence of mosquito-borne viral diseases such as dengue are increasing globally (Pang et al., 2017). Over half of the world’s population are at risk of contracting dengue viruses, with around 105 million people a year estimated to be infected (Bhatt et al., 2013; Cattarino et al., 2020). With conventional arbovirus disease control methods failing to compete with increasing prevalence of such diseases, alternative biocontrol methods have been developed (Bonizzoni et al., 2013; Mayer et al., 2017). Many of these biocontrol methods target the primary vector of these pathogens, the *Aedes aegypti* mosquito. One such method consists of introgressing the bacterial endosymbiont *Wolbachia* into local populations of *Ae. aegypti* mosquitoes (Ross, Turelli, et al., 2019). *Wolbachia* is not naturally present in *Ae. aegypti* (Gerth et al., 2014; Ross, Callahan, et al., 2020), but multiple strains have been stably transinfected into this species (Ant et al., 2018; Fraser et al., 2017, 2020; McMeniman et al., 2009; Walker et al., 2011; Xi et al., 2005). Two strains of *Wolbachia* have undergone field releases as part of *Wolbachia* introgression release strategies in *Ae. aegypti, w*Mel – transinfected from *D. melanogaster* and *w*AlbB – transinfected from *Ae. albopictus* (Ant et al., 2018; Flores et al., 2020; Walker et al., 2011; Xi et al., 2005). These two strains impart two critical features on their host that underpin the biocontrol method. First, they modify host mating outcomes through cytoplasmic incompatibility (CI). In CI, *Wolbachia* infection in males results in the embryonic lethality of offspring when mated with uninfected females. *Wolbachia*-infected females can rescue this lethality, giving them a reproductive advantage (Dobson et al., 2002; Serbus et al., 2008). Combined with the maternal transmission of *Wolbachia*, CI drives *Wolbachia* into uninfected populations. Second, *w*Mel and *w*AlbB infection in *Ae. aegypti* reduces the ability of the mosquitoes to transmit viruses such as dengue, Zika, chikungunya, and yellow fever viruses (Aliota, Peinado, et al., 2016; Aliota, Walker, et al., 2016; Ant et al., 2018; Bian et al., 2010; Dutra et al., 2016; Flores et al., 2020; Rocha et al., 2019; van den Hurk et al., 2012; Walker et al., 2011). In biocontrol methods where *Wolbachia* introgression is used, CI and maternal transmission allow for *Wolbachia* to successfully invade field populations and render them less competent at transmitting arboviruses (O’Neill et al., 2018; Pinto et al., 2021; Ryan et al., 2019; Utarini et al., 2021).

Both *w*Mel and *w*AlbB have generally mild impacts on multiple aspects of *Ae. aegypti* fitness, most notably reducing egg longevity (Allman et al., 2020; Ant et al., 2018; Joubert et al., 2016; Walker et al., 2011). However, on the more moderate side, *w*AlbB infection has been shown to significantly reduce the fertility of females derived from old quiescent eggs (Lau et al., 2021), and while both strains are susceptible to higher temperatures, *w*Mel infection has been shown to have a lower temperature tolerance than *w*AlbB (Ross, Axford, et al., 2020; Ross et al., 2017), both of which could impact their success in the field. *w*Mel-infected *Ae. aegypti* have been released in 14 countries to date (Gesto et al., 2021; Hien et al., 2021; O’Neill et al., 2018; Pocquet et al., 2021; Ryan et al., 2019; Tantowijoyo et al., 2020; Utarini et al., 2021; World Mosquito Program, n.d.). Introgression of *w*Mel into *Ae. aegypti* field populations has led to significant reductions in arbovirus incidence across multiple sites (Indriani et al., 2020; O’Neill et al., 2018; Pinto et al., 2021; Ribeiro Dos Santos et al., 2022; Ryan et al., 2019; Utarini et al., 2021; Ivan Dario Velez et al., 2023). *w*AlbB-infected *Ae. aegypti* have been released in Malaysia by *Wolbachia* Malaysia (Nazni et al., 2019) and are being trialled alongside *w*Mel by the World Mosquito Program (WMP) in Mexico (World Mosquito Program, n.d.).

For both *w*Mel and *w*AlbB strains to perform effectively as a biocontrol method, *w*Mel and *w*AlbB must first successfully introgress to high levels then be maintained at high prevalence in the *Ae. aegypti* population. Typically, *Ae. aegypti* populations are monitored to measure the proportion of the *Ae. aegypti* population that carry a *w*Mel or *w*AlbB infection, and this data is used to inform if introgression is successful and stable. *Wolbachia* prevalence in *Ae. aegypti* populations have been shown to fluctuate when monitored over time (Garcia et al., 2019; Hien et al., 2021; Hoffmann et al., 2014; Nazni et al., 2019; O’Neill et al., 2018; Ryan et al., 2019) but are only rarely lost from a population (Hien et al. 2021). The cause of these fluctuations may be due to invasion of wild-type mosquitoes into release areas, reduced fitness of *Wolbachia*-infected mosquitoes relative to wildtype mosquitoes, or alternatively the loss of *w*Mel or *w*AlbB infection from mosquitoes. Garcia et al. (Garcia et al., 2019) found that *w*Mel prevalence post release in Brazil was impacted due to the loss of pyrethroid resistance in mosquitoes, thus making them less fit compared to wildtype mosquitoes and impacting their ability to introgress into the population. High temperatures have also been shown to impact *w*Mel density and maternal transmission both in the laboratory and in the field (Ross, Axford, et al., 2020; Ross et al., 2017). In an Australian field site, a severe heatwave was associated with decreases in *w*Mel density and prevalence; however, the population recovered to 100% prevalence four months after the heat event (Ross, Axford, et al., 2020). A similar impact of heat was seen at two field sites in Vietnam, where *w*Mel prevalence showed seasonal fluctuations with decreases in prevalence corresponding to increasing temperatures (Hien et al., 2021). However, significant local spatial heterogeneity of *w*Mel prevalence was also observed, suggesting that other factors in addition to heat were likely impacting *w*Mel stability. While some work has been done to investigate the impacts of environmental factors other than temperature fluctuations on *Wolbachia* in *Ae. aegypti* (Ant et al., 2018; Dutra et al., 2015; Endersby-Harshman et al., 2019; Paris et al., 2018; Ross et al., 2016; Ware-Gilmore et al., 2021), this area needs further investigation.

The original *w*Mel transinfection was generated in Australian mosquitoes (Walker et al., 2011). When generating a country-specific release colony of *Ae. aegypti, w*Mel-infected Australian *Ae. aegypti* females are backcrossed for multiple generations to male *Ae. aegypti* from the respective country to localise nuclear genetic background. Since both *Wolbachia* and mitochondria are maternally inherited, this results in a release line carrying a *w*Mel infection, country-specific nuclear genetic background, and the mitochondrial genome of the original Australian founder line. This process provides a unique way to assess *Wolbachia* loss in the field as a mosquito that has lost a *Wolbachia* infection can be distinguished from a wild-type mosquito by the presence of an Australian donor mitochondrial genome and no *Wolbachia* infection. Yeap et al., identified a diagnostic mitochondrial SNP in the COI gene that differentiated *w*Mel-infected mosquitoes with an Australian mitochondrion from local *Ae. aegypti* populations in Vietnam, Indonesia, and Brazil. This SNP has been previously used to detect loss of *w*Mel (i.e. individuals that should have a *w*Mel infection but do not) through the sequencing of COI from field collections and was instrumental in helping to understand the basis of initial challenges to *w*Mel establishment in Brazil (Garcia et al., 2019).

Here, we describe the development of two SNP mitochondrial genotyping (mitotyping) assays and their use to provide a multicountry evaluation of *w*Mel loss from *Ae. aegypti* field populations in Vanuatu, Fiji, Kiribati, Brazil, and Colombia. We then exposed *w*Mel- and *w*AlbB-infected mosquitoes to various field-like conditions to test their effects on maternal transmission. Finally, we manipulated *w*Mel- and *w*AlbB-density levels in *Ae. aegypti* to examine the impact of maternal density on maternal transmission.

## Methods

### Field sites and mosquito collection

#### Field sites

Pacific island releases are described in detail in Simmons et al., (2024). Releases in Suva, Fiji occurred between July 2018 and May 2019 as part of the phase 1 release in Fiji. Releases in Port Villa, Vanuatu occurred between October 2018 and March 2019. Releases in Kiribati occurred in both Betio and Bairiki, South Tarawa, between August 2018 and August 2019. However, due to low sample sizes only mosquito collections from Betio were used in this study.

Colombian releases are described in detail (Iván Darío Velez et al., 2023). Initial releases in Aranjuez, Manrique, Santa Cruz comunas occurred between April 2017 and December 2017 as part of large-scale, phase 1 mosquito releases throughout Bello and Medellín. Additional releases occurred in these areas between April 2018 and March 2019 as part of phase 2 releases. These three release sites were used as ‘intervention’ sites as part of a case-control epidemiological study and are referred to as Aranjuez A, Manrique A, and Santa Cruz comunas, respectively in (Ivan Dario Velez et al., 2023).

Releases for the Brazilian sites of Centro and Maravista in Niterói are described in detail in Pinto et al., (2021), while releases for Universitária in Rio de Janeiro is described in detail in Gesto et al., (2021). Three release periods occurred in Centro (as part of Zone 3 releases) and two release periods occurred in Maravista (as part of Zone 2 releases) between September 2017 and July 2019 (Pinto et al., 2021). Three release periods occurred in Universitária (as part of RJ2 releases) between December 2017 and August 2019 (Gesto et al., 2021).

#### Mosquito Collection

Mosquitoes were collected from field sites using BG Sentinel (BGS) traps (Biogents AG, Germany) or Prokopacks (John W Hock, USA). Mosquitoes were morphologically identified as *Ae. aegypti,* then stored in 80% ethanol until further processing.

#### Species confirmation and Wolbachia detection

qPCR was used to confirm *Ae. aegypti* species identity, as well as *Wolbachia* infection status. Adult mosquitoes were homogenised in extraction buffer (10 mM Tris pH 8.2, 1mM EDTA, 50 mM NaCl, 15 mg/mL proteinase K), then incubated at 56 °C for 5 minutes followed by 98°C for 5 minutes. qPCR was performed on clarified mosquito homogenates using a LightCycler 480 II (Roche) and QuantiNova Probe PCR Master Mix (QIAGEN) except for Brazil samples where GoTaq Probe qPCR Master Mix (Promega) was used, according to the manufacturer’s protocols. Two primer sets were used to amplify DNA to confirm *Ae. aegypti* species identity (specific to the *RpS17* gene) and the presence or absence of a *Wolbachia* infection (specific to the *wsp* gene for *w*Mel). Primers/probes are listed in Supplemental Table 1.

#### Mitochondrial genotyping assays

We developed two assays based on mitochondrial SNPs to distinguish if a mosquito had a mitochondrial genome of the original Australian founder line. The first assay was designed to the cytochrome oxidase I (CO1) gene; as described by Yeap et al., the founder mitochondria differs uniquely at site 531 of the CO1 gene. Each reaction contained forward and reverse primers, a probe specific to the founder mitochondrion (A at site 531), and a probe specific to non-founder mitochondrion (G at site 531) (Supplemental Table 1). This assay was validated against cohorts of wild-type *Ae. aegypti* from Fiji, Vanuatu, Kiribati, and Colombia (Supplementary Table 2). Another unique mitochondrial SNP located in the cytochrome oxidase II (CO2) gene was used that distinguished Brazil mitochondrial genomes from founder genomes. Each reaction contained forward and reverse primers, a probe specific to the founder mitochondrion (A at site 3196), and a probe specific to non-founder mitochondrion (G at site 3196) (Supplemental Table 1). This assay was validated against wild-type *Ae. aegypti* from Brazil (Supplementary Table 2). Using a mosquitoes’ mitochondrial origin in combination with data regarding *w*Mel infection status, mosquitoes can be placed into three categories; *w*Mel-infected progeny of release line, wild-type, or progeny of release line which have lost *w*Mel infection (Table 1).

**Table 1.** Mosquito categorisations following mitotyping and infection status analysis.

|  | Founder mitochondria | Not-Founder mitochondria |
| --- | --- | --- |
| ***w*Mel infected** | Progeny of release line | - |
| **Not *w*Mel infected** | Progeny of release line<br>which has lost *w*Mel<br>infection | Wild-type |

#### Mitotyping

Mosquitoes confirmed to be both *Ae. aegypti* and uninfected with *Wolbachia* were mitotyped to assess origin of mitochondria by qPCR using a LightCycler 480 II (Roche) with QuantiNova Probe PCR Master Mix (QIAGEN) except for Brazil samples where GoTaq Probe qPCR Master Mix (Promega) was used, according to the manufacturer’s protocols. Mosquitoes from Fiji, Vanuatu, Kiribati and Colombia were mitotyped using primers and probes designed to the cytochrome oxidase I (CO1) gene (Supplemental Table 1). The qPCR cycling program consisted of denaturation as 95 °C for 10 minutes, followed by 40 cycles of PCR denaturation at 95 °C for 10 seconds, annealing at 66 °C for 15 seconds, and extension at 72 °C for 1 second with a single acquisition.

Mosquitoes from Brazil were mitotyped using primers and probes designed to the cytochrome oxidase II (CO2) gene (Supplemental Table 1). The qPCR cycling program consisted of denaturation as 95 °C for 10 minutes, followed by 40 cycles of PCR denaturation at 95 °C for 10 seconds, annealing at 60 °C for 15 seconds, and extension at 72 °C for 1 second with a single acquisition.

Mosquitoes are assigned a mitotype by scoring probe fluorescence against internal controls that contain mitochondria corresponding to Australian founder or non-founder mitochondrial SNPs using LightCycler 480 II (Roche) End-Point Genotyping software. Combined with data regarding *w*Mel infection status, mosquitoes were classified into three categories as described in Table 1.

#### Estimating wMel prevalence and wMel loss

*w*Mel prevalence represents the proportion of *Ae. aegypti* mosquitoes in a given population that harbour a *w*Mel infection and is calculated as follows:

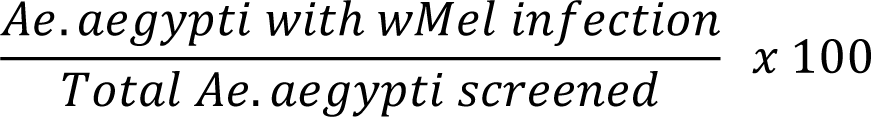

Loss of *w*Mel represents the proportion of the population that, given complete maternal transmission and no loss of *w*Mel, are expected to harbour a *w*Mel infection but do not. As *w*Mel-infected mosquitoes carry an founder mitochondrial genome, this population can be differentiated from wild-type counterparts by the presence of a mitochondrial genotype of the founder release line. Loss of *w*Mel is calculated as follows:

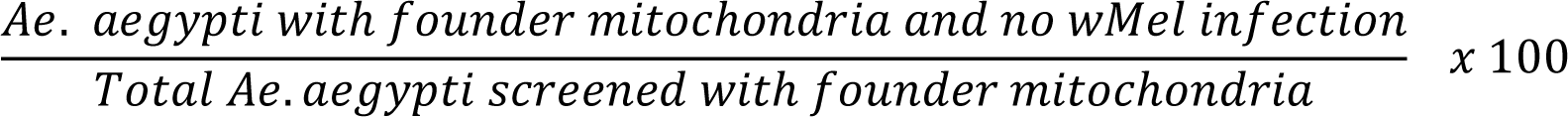

### Laboratory mosquito experiments

#### Mosquito rearing

To better understand what causes loss of *Wolbachia* from individuals, we attempted to recreate conditions in a laboratory setting that we hypothesised could be contributing to loss. Two lines of *Wolbachia* in *Ae. aegypti* were used: *w*Mel- and *w*AlbB-infected *Ae. aegypti* of the same background (Cairns, Australia) as described in Allman et al,. (2023). Both lines have been backcrossed to 10% Cairns wild-type males each generation to reduce the effect of genetic drift of the mosquito lines. For colony maintenance (referred to as standard conditions from herein), eggs were hatched in deoxygenated RO water and thinned into 3L trays of RO water at ∼150 larvae/tray. Larvae were fed Tetra Shrimp Wafers fish pellets (Tetra, Melle, Germany) *ad libitum*. Adults were maintained at a density of ∼450 mosquitoes per cage and maintained at 26 °C and 65% relative humidity (RH) with 12h light:dark cycle and allowed to feed on 10% sucrose *ad libitum*. Female mosquitoes were blood-fed on human volunteers, as approved by the Monash University Human Research Ethics Committee (MUHREC) under Monash University human ethics Project ID: 27690. All adult volunteers provided informed written consent; no child participants were involved in the study.

#### Early blood-feeding

Eggs from *w*Mel- and *w*AlbB-infected *Ae. aegypti* were vacuum hatched in RO water containing Tetra Shrimp Wafers fish food and reared under standard conditions. Pupae were moved to cages and allowed to emerge and collected over 24-hour periods. Females were then blood-fed at the following time points: 0 days post emergence (DPE) (0-24 hours post emergence), 1 DPE (24-48 hours post emergence), 2 DPE (48-72 hours post emergence), and 3 DPE (72-96 hours post emergence). Eggs were collected from each time point and dried. For this process, eggs are collected on filter paper lining a cup filled with RO water. Each filter paper was then placed between layers of paper towel for two hours to remove the majority of water. Papers are then placed on a fresh paper towel, covered with a thin soft-plastic, and left overnight at 26 °C and 65% relative humidity (RH). Filter papers were then stored in containers with saturated potassium chloride to maintain a RH of ∼80% until used. Eggs were then hatched and reared under standard conditions. Adult mosquitoes were collected 3-7 days post emergence and assessed for *Wolbachia* presence by qPCR as described above with *Wolbachia*-specific primers targeting *16S rRNA* gene (Supplementary Table 1).

#### Larval rearing experiments

All larval experiments were performed using standard rearing conditions as described above with the exception of temperature. To mimic temperatures experienced in the three Pacific islands, temperature was controlled over a 24-hour period by maintaining 26 °C for 10 hours, increased over the period of two hours to 31 °C, maintained at 31 °C for 10 hours, then decreased over the period of two hours to 26 °C.

#### Larval starvation

Eggs from *w*Mel- and *w*AlbB-infected *Ae. aegypti* were vacuum hatched in RO water with no food. Two hours post hatching cohorts of 150 1^st^ instar larvae were moved to trays containing three litres of RO water. Larvae were fed fixed amounts of ground Tetra Shrimp Wafers fish food as follows: 0.5 mg/larvae, 1 mg/larvae, 2 mg/larvae, 2.67 mg/larvae (control). Each food amount was fed in total, with no further feedings. Three replicates for each feed condition and two for control conditions were performed for both *w*Mel- and *w*AlbB-infected mosquitoes. Pupae were moved to cages to emerge, where adults were allowed to feed on 10% sucrose *ad libitum*. Adults were blood-fed and eggs were collected and dried as described above, then hatched. Offspring were reared under standard conditions. Adult mosquitoes were collected 3-7 days post emergence and assessed for *Wolbachia* presence by qPCR as described above.

#### Salinity

To simulate brackish water exposure, larvae were reared under two saline conditions where *Ae. aegypti* survival is not substantially impacted (Ramasamy et al., 2014, 2011). Eggs from *w*Mel- and *w*AlbB-infected *Ae. aegypti* were vacuum hatched in RO water containing Tetra Shrimp Wafers fish food. 24-hours post hatching cohorts of 150 1^st^ instar larvae moved to trays containing three litres of RO water containing the following concentrations of Aquarium Salt (API, Mars Fishcare Europe, United Kingdom): 7.5 parts per thousand (ppt), 5 ppt, 2.5 ppt, 0 ppt (control). Larvae were fed Tetra Shrimp Wafers fish food as needed. Three replicates for each Aquarium Salt concentration and two for control conditions were performed for both *w*Mel- and *w*AlbB-infected mosquitoes. Pupae were moved to cages to emerge in RO water with matching aquarium salt concentrations, where adults were allowed to feed on 10% sucrose *ad libitum*. Adults were blood-fed and eggs were collected and dried as described above, then hatched. Offspring were reared under standard conditions (no aquarium salt). Adult mosquitoes were collected 3-7 days post emergence and assessed for *Wolbachia* presence by qPCR as described above.

#### Antibiotic manipulation of wMel and wAlbB density

Eggs from *w*Mel- and *w*AlbB-infected *Ae. aegypti* were vacuum hatched in RO water containing Tetra Shrimp Wafers fish food. 24-hours post hatching, cohorts of 150 1^st^ instar larvae were moved to trays containing three litres of RO water. Larvae were fed Tetra Shrimp Wafers fish food as needed. Pupae were moved to cages to emerge. Each cage contained a 10% sucrose solution with the tetracycline hydrochloride (Sigma-Aldrich) at one of the following concentrations: 0.8 mg/mL, 0.4 mg/mL, 0.2 mg/mL, 0.1 mg/mL, 0.05 mg/mL, or 0 mg/mL (control). The tetracycline-spiked sucrose was refreshed every 2-3 days over a 12-day period. Female mosquitoes were then blood-fed, and tetracycline-spiked sucrose was replaced with sucrose only. Female mosquitoes were then isofemaled and collected 3 days post-blood feeding, and eggs for each individual female collected and hatched without drying. Offspring were reared under standard conditions and adults collected 3-7 days post emergence. Maternal female mosquitoes were homogenised in buffer ATL and proteinase K (1 mg/mL) and incubated at 56 °C and 96 °C for 5 minutes each, before total DNA was isolated from individual tissues using the QIAamp 96 DNA QIAcube HT kit (Qiagen) following manufacturer’s instructions. Offspring mosquitoes were homogenised in extraction buffer and incubated at 56 °C and 96 °C for 5 minutes each and clarified mosquito homogenates used for qPCR. Relative *Wolbachia* density was determined in each individual mosquito using qPCR with primers to amplify a fragment of the *Wolbachia 16S rRNA gene*, and the reference *Ae. aegypti RpS17* gene (Supplementary Table 1). qPCR was performed as described above. *Wolbachia* density was quantified relative to *rps17* using the ΔCT method (2^CT^(reference)/2^CT^(target)) (Pfaffl, 2001).

#### Statistical analysis

All data collected were analysed using GraphPad Prism v9 or R version 4.1.1. The Wilson/Brown method was performed to assess 95% confidence intervals using the technical replicates. Correlations were calculated using Kendall’s rank correlation in R.

## Results

### *Loss of* wMel *in field mosquitoes varies substantially between field sites*

Current WMP release strategies result in *w*Mel-infected mosquitoes that have a mitochondrial genotype associated with the Australian founder female mosquito. We used this to our advantage to better understand *w*Mel introgression dynamics. We surveyed mosquitoes from five different countries across multiple time points to gain a comprehensive understanding of how frequently *w*Mel is lost from mosquito populations in the field. We assessed mosquitoes to confirm their species identity, *Wolbachia* infection status, and mitotype (founder or non-founder). Here, we report on two population statistics: *w*Mel prevalence and loss of *w*Mel (see methods). While we present *w*Mel loss as point estimates of the population proportion to have lost *w*Mel, it should be noted that this is reflective of cumulative *w*Mel loss in the population as a whole over time, rather than a measure of *w*Mel loss in the current generation. This is because the loss of *w*Mel event may have occurred at any time in the maternal lineage of the individuals tested and has not necessarily been lost in the lifetime of the analysed individual.

### Pacific Island Countries

We first measured *w*Mel loss in three release sites in Pacific Island countries (Suva, Fiji; Port Villa, Vanuatu; Betio, South Tarawa, Kiribati) across multiple timepoints (Figure 1). With the exception of Suva, Fiji, populations were analysed after releases ceased. In Suva, subsequent short periods of releases continued after initial mitotyping analysis in sites close to where sampling was performed; however, samples for mitotyping were taken from sites where releases had ceased. In Port Villa, Vanuatu, *w*Mel prevalence ranged between 68.1 – 71% across three time points spanning a year (Figure 2A). In the same populations, loss of *w*Mel was calculated to be between 1.8 – 3%. Similarly, *w*Mel prevalence in Suva, Fiji, ranged between 57.4 – 83.6% across five time points spanning one year three months, during which time loss of *w*Mel ranged between 0 – 3.9% (Figure 2B). The release site of Betio in South Tarawa, Kiribati, experienced *w*Mel prevalence similar to that of Vanuatu and Fiji, ranging between 52.3 – 80.5% across four time points spanning seven months (Figure 2C). However, loss of *w*Mel during this period was higher in Kiribati than on the other two Pacific countries, ranging between 2.6 – 12.8%. Data from both Fiji and Kiribati suggested that loss of *w*Mel may have contributed to the reduced prevalence of *w*Mel. For example, the highest level of *w*Mel loss in Fiji occurs during the same time *w*Mel prevalence dips in mid 2019. Similarly, *w*Mel prevalence in Kiribati increased substantially as loss of *w*Mel decreases.

**Figure 1.**
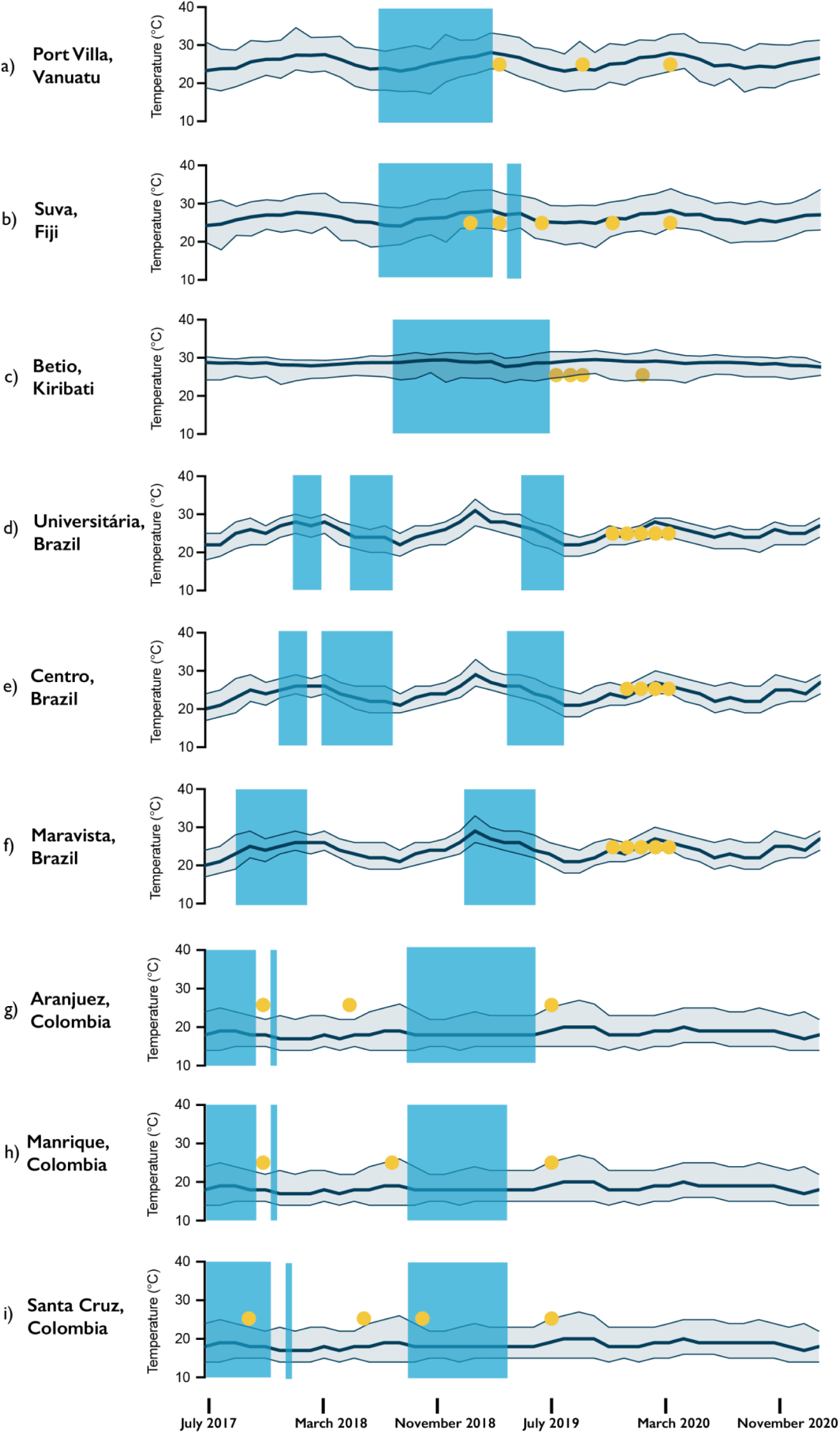
Timeline of *Wolbachia*-infected *Ae. aegypti* release periods, mitotyping collection timepoints, and ambient temperatures in surveyed areas. Blue rectangles represent periods where *Wolbachia*-infected *Ae. aegypti* were released in indicated field sites. Yellow circles represent timepoints where mosquitoes were collected from the indicated field sites and assessed for mitotyping purposes. a) - c) show mean, minimum, and maximum temperatures of Vanuatu, Fiji, and Kiribati respectively, during the described period as reported by http://www.bom.gov.au/. d) - f) show mean, minimum, and maximum temperatures of Rio de Janeiro, Brazil, during the described period as reported by worldweatheronline.com. g) - i) show mean, minimum, and maximum temperatures of Medellin, Colombia, during the described period as reported by worldweatheronline.com.

**Figure 2.**
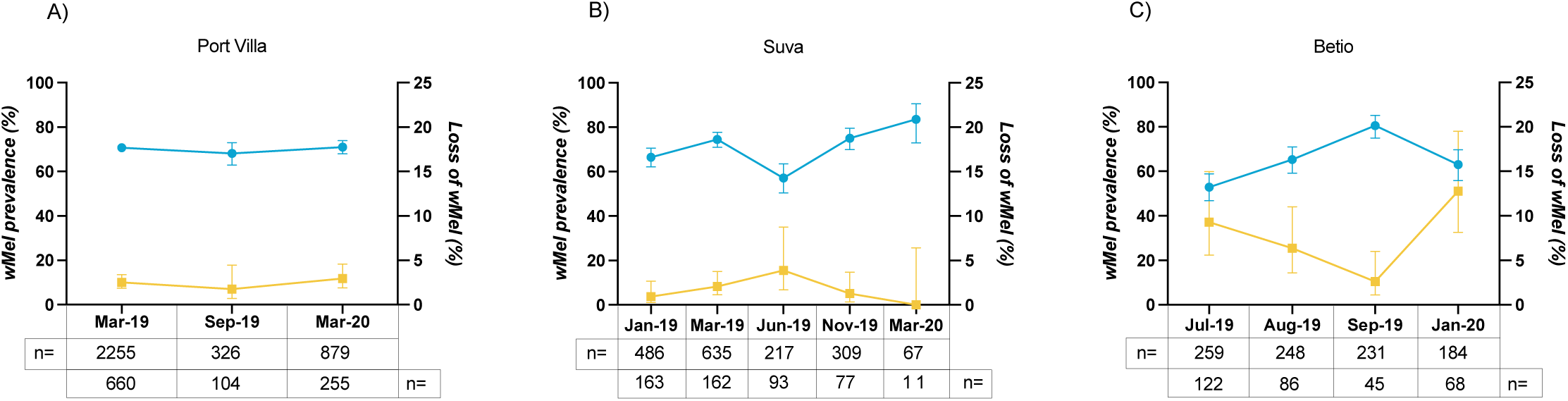
*Wolbachia* prevalence and *Wolbachia* loss vary over time in Pacific Island Countries. Graphs show *w*Mel prevalence (left axis) represented by blue lines and dots, and loss of *w*Mel (right axis) represented by yellow lines and dots over time in *Ae. aegypti* populations in Port Villa, Vanuatu (A), Suva, Fiji (B), Betio, Kiribati (C). Error bars represent confidence intervals. Sample sizes depicting number of mosquitoes assayed to estimate *w*Mel prevalence for each time point are depicted in the top n= line under respective dates. Sample sizes depicting individuals without a *w*Mel infection that were assessed for loss of *w*Mel are depicted in the bottom n= line under respective dates.

Previous studies have shown *w*Mel is sensitive to high temperatures, with decreased density and loss of *w*Mel reported after heat exposure (Ross, Axford, et al., 2020; Ross et al., 2017). We therefore compared the ambient temperature ranges of the three Pacific Island countries over the period studied (Figure 1A-C). Mean ambient air temperatures reported during the monitored period in Vanuatu ranged between 23.4 – 27.9°C, with the highest recorded temperature at 33°C. Similarly, mean ambient temperatures reported in Fiji ranged between 24.8 – 28.2°C, with the highest recorded temperature during monitoring being 33.9°C. Lastly, mean ambient air temperatures reported during the monitored period in Kiribati ranged between 28.9– 29.5°C, with the highest recorded temperature at 32.2°C (Figure 1). These data suggest that temperature differences between the Pacific Islands is unlikely to be a major contributing factor to differences observed in *Wolbachia* loss as decreases in *w*Mel prevalence and/or loss are not associated with high temperature periods.

### Brazil

Next, we compared three release sites within Brazil across multiple time points (Figure 1). The first site, Universitária, is located in Rio de Janeiro, while the second and third sites, Centro and Maravista, are located in Niterói. Universitária had *w*Mel prevalence that decreased steadily over five months from 54.4% to 35.2% (Figure 3A). During this period, loss of *w*Mel varied between 6 – 8.8% remaining consistent over the sampling period. *w*Mel prevalence in the release area of Centro decreased from 43.3% to 17.2% between December 2019 and January 2020, then was measured to be 19.2% and 25.35% over the following two months (Figure 3B). Interestingly, between December 2019 and January 2020 loss of *w*Mel increased from 3.7% to 20.4%. During the following two months, loss of *w*Mel measured at 18.2% and 10% respectively. Finally, across a period of five months, the release site of Maravista had a *w*Mel prevalence ranging between 27% and 41.4% (Figure 3C). During this time, loss of *w*Mel increased from 0% to 16.7%.

**Figure 3.**
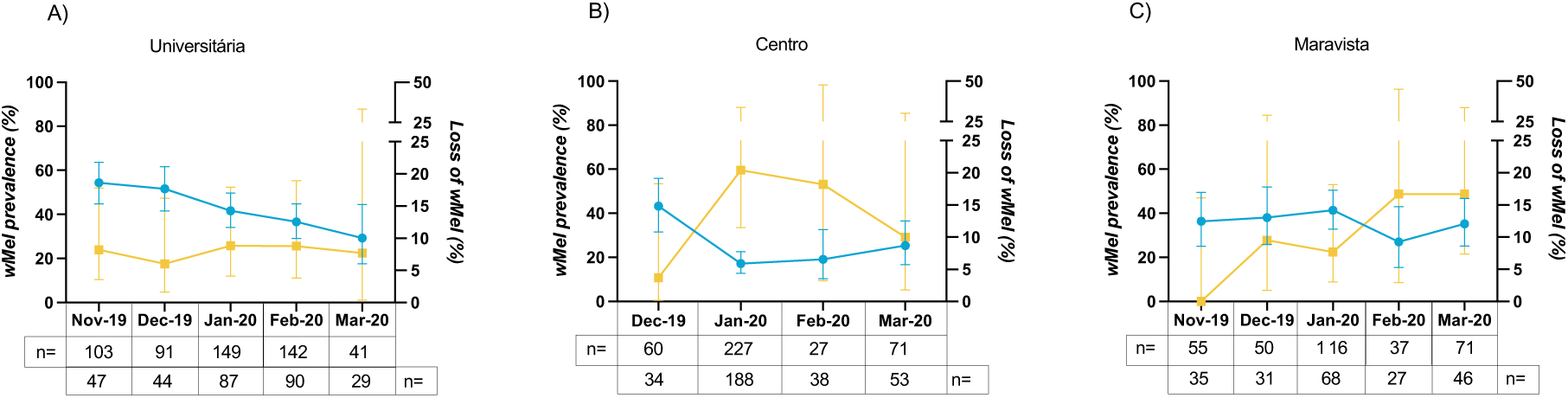
*Wolbachia* prevalence and *Wolbachia* loss vary over time in Brazil. Graphs show *w*Mel prevalence (left axis) represented by blue lines and dots, and loss of *w*Mel (right axis) represented by yellow lines and dots over time in *Ae. aegypti* populations in Universitária, Rio de Janeiro, Brazil (A), Centro, Niterói, Brazil (B), Maravista, Niterói, Brazil (C). Error bars represent confidence intervals. Sample sizes depicting number of mosquitoes assayed to estimate *w*Mel prevalence for each time point are depicted in the top n= line under respective dates. Sample sizes depicting individuals without a *w*Mel infection that were assessed for loss of *w*Mel are depicted in the bottom n= line under respective dates.

During the monitored period, mean ambient air temperatures in Niterói and Rio de Janeiro, Brazil, were reported to range between 24 – 28°C with the highest recorded ambient air temperature during this period being 30°C (Figure 1D-F). Despite these three sites being reasonably close geographically and experiencing similar temperatures, we find that *w*Mel loss is associated with decreases in *w*Mel prevalence at varying amounts across the different sites and time periods and does not necessarily occur during periods of high temperature.

### Colombia

We then compared three release sites within the city of Medellín, Colombia that span an area <5km, across multiple time points (Figure 1). These sites differ from those previously discussed as releases of *w*Mel-infected *Ae. aegypti* were ongoing in between the sampling periods of this study. Each of these three sites, Aranjuez, Manrique, and Santa Cruz, subsequently show varied levels of *w*Mel population prevalence, influenced in part by the *w*Mel releases. However, each of the three release sites show low *w*Mel population prevalence, below 30%, during at least one of the studied time points (Figure 4). Surprisingly, however, very little or no loss of *w*Mel was seen in these three sites. In Aranjuez, loss of *w*Mel was identified at 8.3% at one time point, however no identification of mosquitoes that had lost a *w*Mel infection were identified at any other time points (Figure 4A). Similarly, no loss of *w*Mel was identified at either Manrique or Santa Cruz across the studied time points, despite the occasional low population prevalence of *w*Mel (Figure 4B-C).

**Figure 4.**
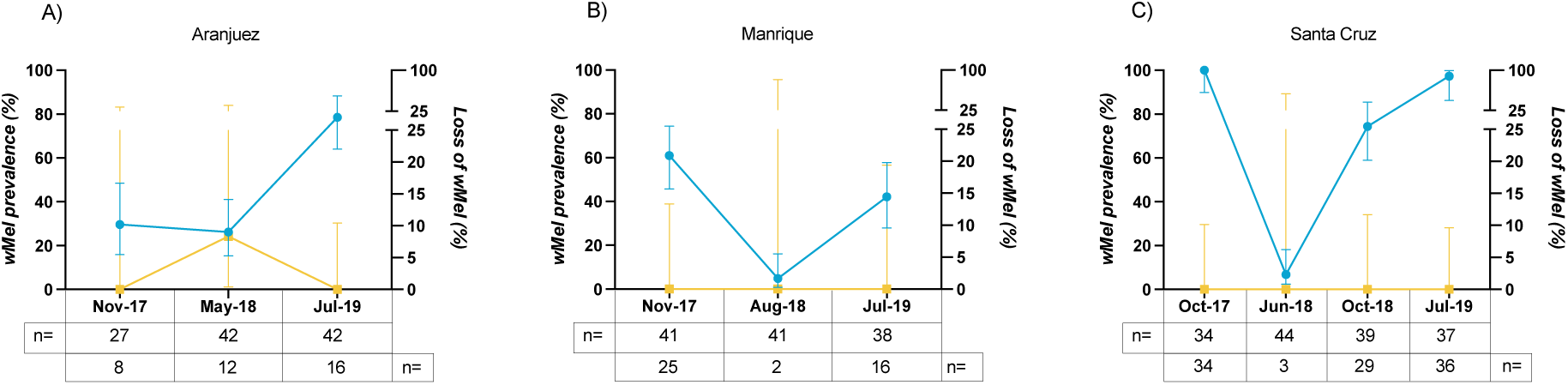
*Wolbachia* prevalence and *Wolbachia* loss over time in Colombia. Graphs show *w*Mel prevalence (left axis) represented by blue lines and dots, and loss of *w*Mel (right axis) represented by yellow lines and dots over time in *Ae. aegypti* populations in Aranjuez, Medellín, Colombia (A), Manrique, Medellín, Colombia (B), Santa Cruz, Medellín, Colombia (C). Error bars represent confidence intervals. Sample sizes depicting number of mosquitoes assayed to estimate *w*Mel prevalence for each time point are depicted in the top n= line under respective dates. Sample sizes depicting individuals without a *w*Mel infection that were assessed for loss of *w*Mel are depicted in the bottom n= line under respective dates.

#### Laboratory simulation of environmental parameters does not result in Wolbachia loss

High ambient temperatures are known to impact *w*Mel prevalence in field populations (Hien et al., 2021; Ross, Axford, et al., 2020). However, our data, particularly from the Pacific Islands, suggest that other environmental conditions that mosquitoes are exposed to may also affect both *Wolbachia* stability in *Ae. aegypti.* Therefore, we investigated the effects of potential field and environmental conditions other than high temperature on the maternal transmission of both *w*Mel and *w*AlbB.

### Impact of early blood feeding on maternal transmission of both wMel and wAlbB

Due to reduced feeding rates of younger *Ae. aegypti* mosquitoes (Alto et al., 2003), *Wolbachia* maternal transmission assays are generally performed on mosquitoes that are 5 days old or older (Ant et al., 2018; Axford et al., 2016; Fraser et al., 2017). At this age *Wolbachia* densities in ovaries are much higher than in newly emerged adult females. To understand if lower *Wolbachia* density in young females impacted maternal transmission, we blood-fed *w*Mel- and *w*AlbB-infected *Ae. aegypti* at zero, one, two, and three days post-emergence, then measured the maternal transmission of *w*Mel and *w*AlbB to offspring. *w*Mel- and *w*AlbB-infected *Ae. aegypti* blood-fed at zero days post-emergence produced only 21 and 1 offspring respectively. Maternal transmission to each of these cohorts was 100% with 95% confidence intervals (CIs) of 84.5-100 and 5.1-100, respectively, due to low sample size. *w*Mel- and *w*AlbB-infected *Ae. aegypti* blood-fed at one, two, and three days post emergence all experienced complete maternal transmission of *Wolbachia* to (95% CI 96.1-100) (Table 2) suggesting that lower *Wolbachia* density in young mothers does not have an impact on maternal transmission of *Wolbachia*.

**Table 2:** Maternal transmission of *Wolbachia* in *Ae. aegypti* post-exposure. DPE – days post emergence, ppt – parts per thousand.

| Condition | Strain | Treatment | Maternal Transmission %<br>(+/- 95% CI) | Number of progeny tested |
| --- | --- | --- | --- | --- |
| Early Blood-feed | wMel | Blood fed at 0 DPE | 100 (84.5 – 100) | 21 |
|  |  | Blood fed at 1 DPE | 100 (96.1 – 100) | 94 |
|  |  | Blood fed at 2 DPE | 100 (96.1 – 100) | 94 |
|  |  | Blood fed at 3 DPE | 100 (96.1 – 100) | 94 |
|  | wAlb | Blood fed at 0 DPE | 100 (5.1 – 100) | 1 |
|  |  | Blood fed at 1 DPE | 100 (96.1 – 100) | 94 |
|  |  | Blood fed at 2 DPE | 100 (96.1 – 100) | 94 |
|  |  | Blood fed at 3 DPE | 100 (96.1 – 100) | 94 |
| Starvation | wMel | 4 mg/larvae - Control | 100 (92.6 – 100) | 48 |
|  |  | 2 mg/larvae | 100 (94.9 – 100) | 72 |
|  |  | 1 mg/larvae | 100 (94.9 – 100) | 72 |
|  |  | 0.5 mg/larvae | 100 (92.6 – 100) | 48 |
|  | wAlb | 4 mg/larvae - Control | 100 (92.6 – 100) | 48 |
|  |  | 2 mg/larvae | 100 (94.9 – 100) | 72 |
|  |  | 1 mg/larvae | 100 (93.5 – 100) | 55 |
|  |  | 0.5 mg/larvae | - | 0 |
| Salinity | wMel | 0 ppt - Control | 100 (92.6 – 100) | 48 |
|  |  | 5 ppt | 100 (94.9 – 100) | 72 |
|  |  | 7.5 ppt | 100 (94.9 – 100) | 72 |
|  | wAlb | 0 ppt - Control | 100 (92.6 – 100) | 48 |
|  |  | 5 ppt | 100 (94.9 – 100) | 72 |
|  |  | 7.5 ppt | 100 (94.9 – 100) | 72 |

### Impact of salinity and starvation on maternal transmission of Wolbachia

We tested the maternal transmission of *w*Mel- and *w*AlbB-infected *Ae. aegypti* when larvae were exposed to either saline water conditions or starvation. To mimic conditions exposed to in the field, mosquitoes were reared during all life stages using a cycling temperature range from 26°C – 31°C, as these temperatures simulate typical daily temperature cycles experienced in the three Pacific sites studied above (Figure 1A-C).

To mimic brackish water salt concentrations where *Ae. aegypti* survival is not substantially impacted (Ramasamy et al., 2014, 2011) larvae were reared under two saline conditions of 5 and 7.5 ppt. Under these concentrations, maternal transmission of *w*Mel and *w*AlbB from mothers to offspring was 100% (95% CI 92.6-94.9 – 100) (Table 2). In a separate experiment, larvae were starved by decreasing standard lab food amounts down to ½, ¼ or 1/8 of standard food allotment. Under the lowest food diets, larvae were delayed in development and had decreased pupation rates as noted in the decreased sample size. In particular, no adults emerged from *w*AlbB larvae fed on the lowest food diets. Again, when larvae were reared under these starvation conditions, maternal transmission of *w*Mel and *w*AlbB from mothers to offspring was 100% (95% CI 92.6-94.9 – 100) (Table 2). Biological replicates were performed for each of these assays, with maternal transmission of both *w*Mel and *w*AlbB again found to be 100% for each condition, while *w*AlbB larvae did not survive to adulthood under the most nutritionally deficient condition (Supplemental Table 3).

#### Importance of maternal Wolbachia density on maternal transmission of Wolbachia

Lastly, we wanted to understand the importance of maternal *Wolbachia* density for maternal transmission of *Wolbachia*. To do this, we manipulated maternal *w*Mel and *w*AlbB density with tetracycline and collected each individual mothers’ offspring. Then we measured the amount of *w*Mel and *w*AlbB present in each mother as well as the amount of *w*Mel and *w*AlbB present in a subset of her offspring. Maternal females infected with *w*Mel were measured to have densities ranging between 7.13 to 0.0136 *w*Mel per host cell (Figure 5A). Maternal transmission of *w*Mel was 100% until the maternal density of *w*Mel reached 0.426 *w*Mel per host cell, where we observed the first instance of incomplete maternal transmission. Interestingly, incomplete maternal transmission was not guaranteed below this *w*Mel density level but was seen sporadically throughout individuals with density lower than 0.426 *w*Mel per host cell. This is likely a result of the time lag between when the mother laid her eggs and when she was sampled for measuring *Wolbachia* density during which she was not exposed to antibiotics. The magnitude of maternal transmission perturbation experienced by affected individuals was overall quite small. The greatest loss of maternal transmission occurred in an individual where 83.33% of her offspring still had a *w*Mel infection. Complete maternal transmission was also seen in individuals with very low *w*Mel density. The impacts of maternal tetracycline treatment were more strongly observed in the density of *w*Mel in offspring, where we observed a corresponding decrease in the *w*Mel density as maternal density decreased. Maternal *w*Mel density was positively correlated to mean offspring density (Kendall’s tau=0.541, P=8.71 x 10^-13^).

**Figure 5.**
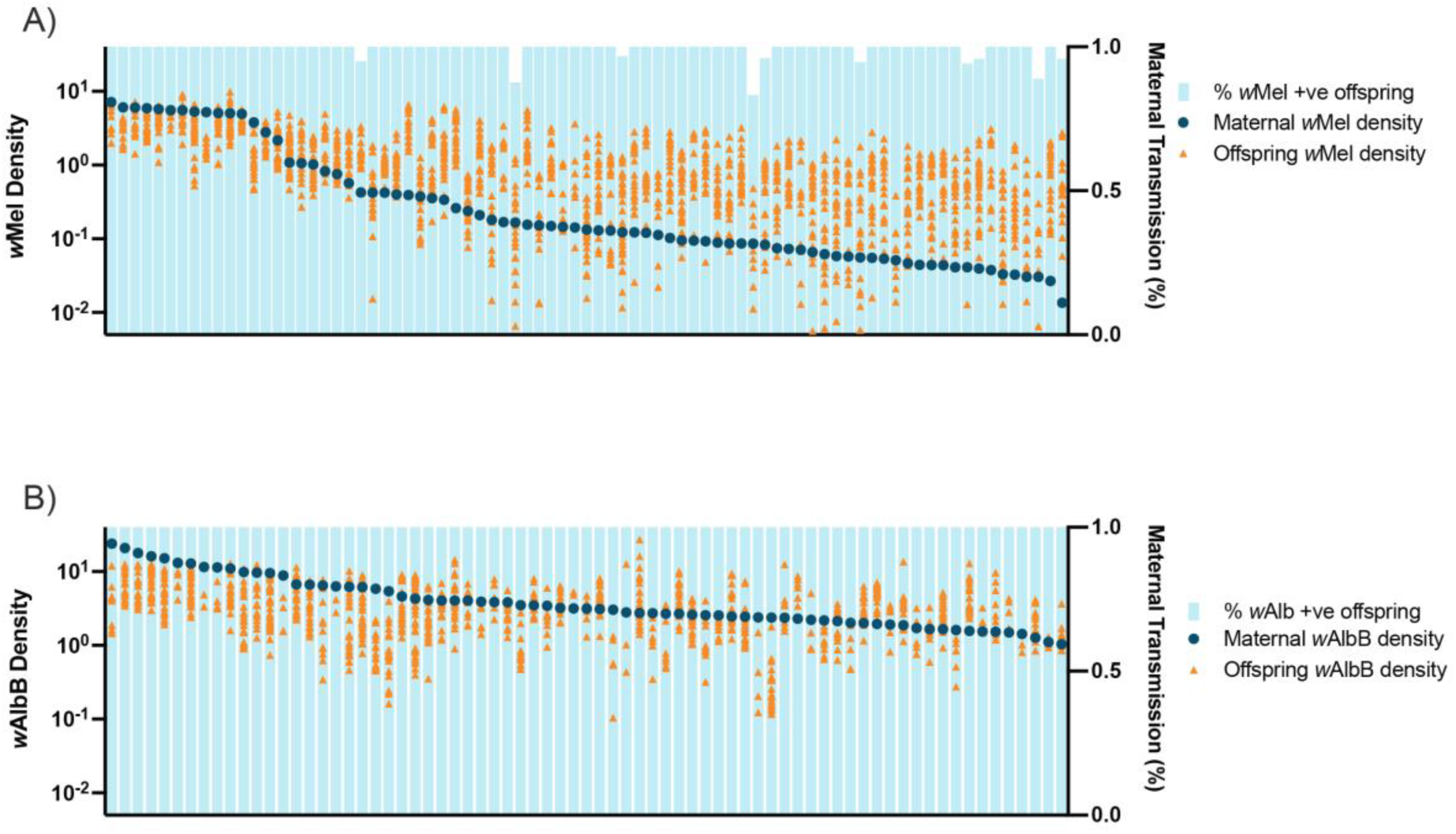
Maternal *Wolbachia* density has minimal impacts on its maternal transmission but does impact *Wolbachia* density in offspring. Graph A) represents *Ae. aegypti* mothers and offspring infected with *w*Mel. Graph B) represents *Ae. aegypti* mothers and offspring infected with *w*AlbB. In both graphs, data points are stratified by maternal *Wolbachia* density which are represented as large blue dots (left axis), orange triangles represent density of each respective mothers’ offspring (left axis), and the light blue foreground bars represent the percentage of each mothers’ offspring that carry a *Wolbachia* infection (right axis).

Maternal females infected with *w*AlbB were measured to have densities ranging between 23.9 to 1.03 *w*AlbB per host cell (Figure 5B), much higher than *w*Mel overall. Maternal transmission was found to be 100% for each *w*AlbB-infected individual at these densities. Maternal *w*AlbB density was also positively correlated to mean offspring density (Kendall’s tau=0.365, P=5.06 x 10^-6^). The difference in maternal transmission between *w*Mel and *w*AlbB is likely due to the higher starting density for *w*AlbB. It should be noted, however, that *w*Mel density is much higher in the ovaries than *w*AlbB (Fraser 2020). As ovarian density is a more accurate measure of the *Wolbachia* that can be maternally transmitted, it is possible that *w*Mel initial densities could be much higher than *w*AlbB in ovaries as we measured whole body density.

## Discussion

*Wolbachia* introgression methods are currently being implemented globally in 15 countries to reduce the transmission of medically important viruses (Nazni et al., 2019; World Mosquito Program, n.d.). The goal of our study was to determine to what extent *w*Mel loss may be playing a role in the variation in *Wolbachia* prevalence observed in global field sites and investigate its potential causes. We made estimates of the fidelity of *w*Mel maternal transmission by measurement of mitotypes and *w*Mel infection status. Interestingly, we found every field site experienced some loss of *w*Mel which varied greatly between field sites. Encouragingly, *w*Mel remained established in each field sites with higher levels of loss. We then attempted to identify factors in laboratory conditions that may undermine maternal transmission and found that early blood feeding, increased saline conditions, and starvation did not individually impact maternal *w*Mel or *w*AlbB transmission when replicated in a lab setting. Finally, we artificially altered maternal *w*Mel and *w*AlbB densities and measured maternal transmission to understand the interplay between maternal density and transmission. We found that despite significant reduction of *w*Mel and *w*AlbB in the mother, maternal transmission was nearly complete in both *w*Mel and *w*AlbB mosquitoes.

While our data suggest *w*Mel loss is contributing to some of the variation in *w*Mel levels observed in field sites, it also suggests that other factors are contributing as well. For example, in some Brazilian sites we observe sustained low levels of *w*Mel which can influence introgression in multiple ways. The success of *w*Mel introgression is driven by the beneficial effects of CI and maternal transmission which require substantial *w*Mel levels (modelling predicts >30%) to effectively drive *Wolbachia* into populations (Turelli & Barton, 2017). These beneficial effects also offset fitness costs incurred by *w*Mel infection. However, when *w*Mel prevalence and CI levels are low, the fitness benefit to *w*Mel-infected females is also significantly reduced likely resulting in a reduced relative fitness compared to wild-type females, ultimately further impacting the ability of *w*Mel to introgress into populations.

Temperature has been shown to influence *w*Mel prevalence in *Ae. aegypti* field populations (Hien et al., 2021; Ross, Axford, et al., 2020). However, the sites studied here show no great variation in ambient temperatures despite displaying varying levels of *w*Mel loss. In particular, we observe similar differences between mean and max ambient air temperatures in all three Pacific sites during both times of high (July 2019) and low (September 2019) loss of *w*Mel in Kiribati. Breeding container type, size, and exposure to sun have been shown to impact temperatures of breeding water and thus *w*Mel levels and maternal transmission (Hien et al., 2021; Ross, Ritchie, et al., 2019). No studies have been performed to assess the types, frequency, and sun exposure of breeding containers and whether this varies across the Pacific Islands. Therefore, we cannot eliminate the possibility that substantial differences in breeding water temperatures may underlie the differences we observe in *w*Mel loss. However, our data indicates there may be other environmental factors impacting loss of *w*Mel.

Multiple mathematical models have been used to predict *Wolbachia* population dynamics and/or epidemiological impact. The models are informed by critical traits conferred by *w*Mel such as strength of cytoplasmic incompatibility and maternal transmission to model invasion dynamics, as well as strength of virus inhibition to model impacts on dengue transmission. While most models incorporate some measure of the fitness cost of *Wolbachia* infection, they often assume complete CI and maternal transmission (Ferguson et al., 2015; Hughes & Britton, 2013; Walker et al., 2011). Even those models that do incorporate incomplete maternal transmission assume only a reduction of 5-10% (Adekunle et al., 2019; Dorigatti et al., 2018). Our data reported here, suggests that incorporating higher levels of incomplete maternal transmission into models of *Wolbachia* population dynamics and epidemiological impact may be warranted to more accurately determine *Wolbachia* spread and impact especially with more extreme environments occurring as a result of climate change.

There are multiple factors that feed into the field data presented here which can impact our overall assessment and interpretation. The data we presented here cover a 5–12-month snapshot of *Wolbachia* prevalence and loss at each release site with 3-6 sampling periods and in some cases low numbers of *Wolbachia*-negative mosquitoes. To have a more comprehensive view, we would ideally monitor loss of *Wolbachia* across longer timeframes to better understand the impacts of seasonality and extreme weather events. Additionally, increased sampling temporally as well as sampling points per collection would allow for increased accuracy in our measurements of *Wolbachia* loss and may give insights into factors which effect this. Our analysis here also assumes that the only way to obtain a *Wolbachia*-negative mosquito with an Australian mitochondrial genotype is through *Wolbachia* loss. Paternal transmission of mitochondria, combined with stochastic loss of one mitotype or incomplete CI (H. L. Yeap et al., 2015), can result in this genotype. However multiple studies suggest paternal transmission events are very rare even on their own, and thus we feel our assumption is justified (Nunes et al., 2013; H. L. Yeap et al., 2015). Despite these limitations, these data provide significant insight into the contribution of *Wolbachia* loss to *Wolbachia* fluctuations observed in the field and highlight the need to better understand other factors that could delay or prevent *Wolbachia* establishment and stability in the field.

In an effort to identify novel environmental factors that, in combination with field-like temperatures, could impact *Wolbachia* stability, we exposed developing *w*Mel and *w*AlbB infected larvae to saline water and starvation conditions to understand if this would impact the maternal transmission of *Wolbachia*. Although *Ae. aegypti* typically breed in fresh water, in some near-coastal sites where *w*Mel-infected *Ae. aegypti* are released (e.g. Kiribati, Vanuatu, Universitária, Centro, Maravista), mosquitoes may breed in brackish water sites (Jude et al., 2012; Ramasamy et al., 2011; Yee et al., 2014). Food sources may also be limited in some breeding sites, leading larvae to experience starvation during growth. Here, when *Wolbachia*-infected *Ae. aegypti* hosts were exposed to both saline water conditions or starvation conditions, maternal transmission remained complete. This indicates that field-like temperature conditions and individual exposure to saline water and starvation conditions does not impact the maternal transmission of *Wolbachia* in *Ae. aegypti.* Similarly, taking an early blood-meal did not cause perturbation to maternal transmission of *w*Mel or *w*AlbB, indicating it is unlikely to be impacting field loss of *Wolbachia.* However, given these conditions could occur simultaneously in field conditions, further work could look to assess the impact of these conditions in combination as well as across multiple generations as small impacts may be compounded over time. Identifying these influencing factors may allow for implemental avoidance of these factors, or for *w*Mel or *w*AlbB to be selected to be less resistant to these factors.

While starvation conditions did not impact maternal transmission of either *Wolbachia* strain, we did observe somewhat different impacts on larval survival. In both experimental repeats, *w*AlbB larvae did not survive to adulthood when reared under the most nutrient deficient conditions. On the other hand, *w*Mel had reduced survival in one repeat (Table 2), and no survival to adulthood in a second repeat (Supplemental Table 3), suggesting that perhaps *w*AlbB may be more costly to larvae under more nutrient-deficient conditions. Our data are consistent with a previous study indicating a higher fitness cost of *w*AlbB compared to wildtype and *w*Mel-infected larvae. When larvae were starved at the third-instar stage, *w*AlbB-infected larvae showed decreased survival compared to *w*Mel and wildtype larvae (Ross et al., 2016). Only one other study has looked at survival to adulthood when larvae were fed under low nutrition conditions starting at the first instar stage. They found *w*AlbB-infected larvae had no significant reduction in survival compared to wildtype larvae which showed >93% survival (Axford et al., 2016). However, the low nutrition condition used in this study used approximately 2-2.5 times the amount of food per larvae as used in our study. Additionally, our study also included temperature ranges to mimic those observed in the Pacific Islands, which may have exacerbated the impacts of starvation alone.

Natural variation in *Wolbachia* density is not very large among *Ae. aegypti* mosquitoes (Amuzu & McGraw, 2016); therefore, we artificially modulated it using antibiotics allowing us to measure its impact on maternal transmission. Surprisingly, we found that disrupting maternal transmission of *Wolbachia*, by modulating maternal *Wolbachia* densities with antibiotics, was very difficult to achieve. Even when *w*Mel density was significantly reduced, by almost two logs, maternal transmission was not necessarily incomplete. Instead, our data showed that maternal density had more of an impact on mean offspring than *Wolbachia* density. However, it is worth noting that we are measuring *w*Mel DNA levels which may not be representative of viable *w*Mel. Additionally, the measurement of maternal density here is also limited by the delay between egg lay and the collection. Maternal densities were influenced by active antibiotic treatment prior to oviposition, and mothers were not collected for density measurement until after oviposition. Therefore, the densities measured here may be lower than they would be at the time of oviposition.

Here, we present a multi-country assessment of *Wolbachia* loss from field *Ae. aegypti*. We found *Wolbachia* loss in *Ae. aegypti* to vary between release sites, and to be a significant contributing factor to fluctuations of *Wolbachia* prevalence in some field sites. We also show when individually tested, the environmental stressors used here have no impact on maternal transmission. Given these conditions may occur in tandem in field sites, there is further scope to understand the effect of compounding environmental conditions on the maternal transmission of *Wolbachia* in *Ae. aegypti.* We show the importance of understanding levels of *Wolbachia* loss in *Ae. aegypti* populations and highlight the need for further work in establishing causes of *Wolbachia* loss as well as incorporating field estimates of *Wolbachia* loss in modelling.

## Acknowledgements

We would like to acknowledge the efforts of the Pacific Island implementation teams for their efforts in mosquito collections, and the efforts of Ritzel Gimeno and Limom Lim for technical assistance.

## Supporting Information

**Supplementary Table 1.**
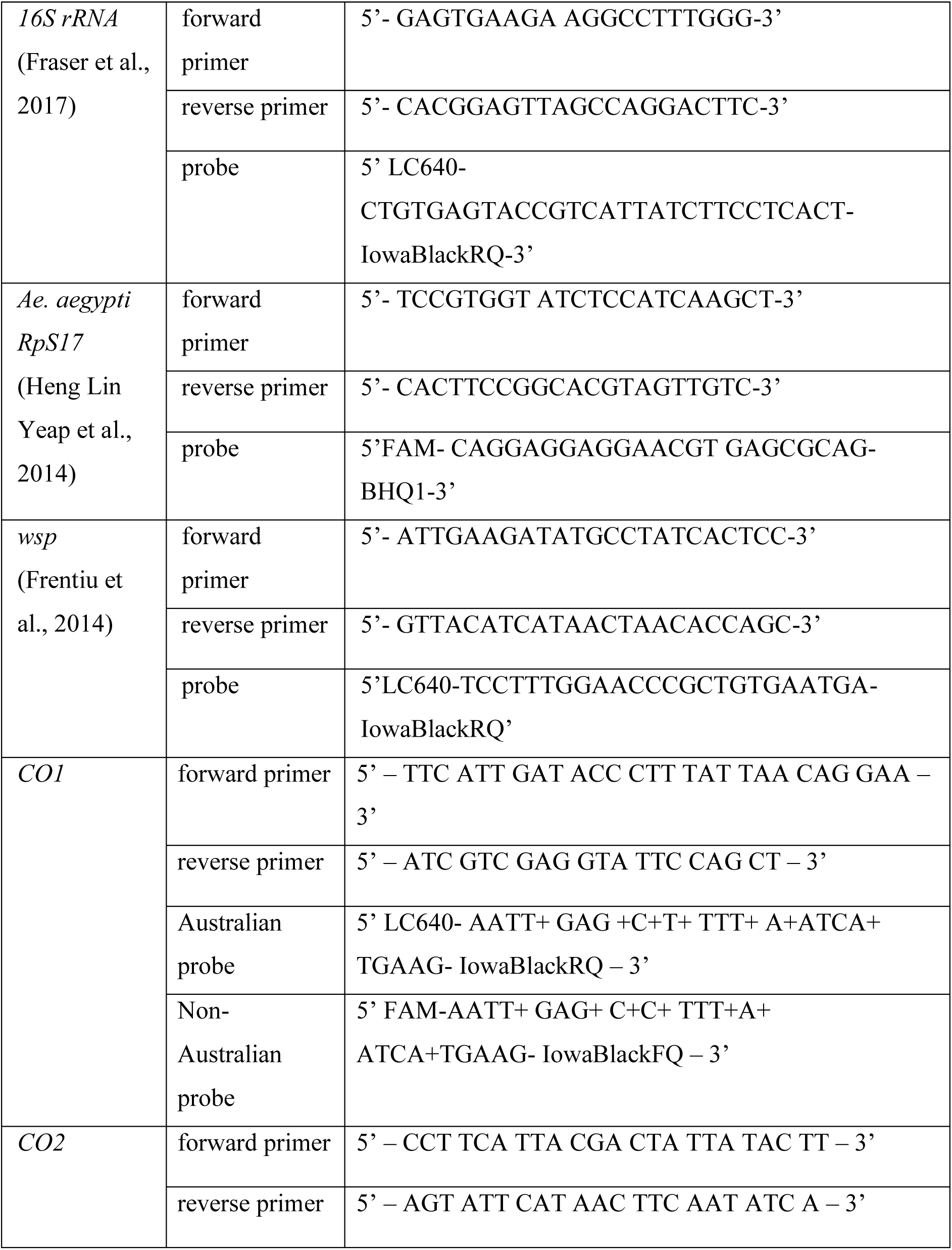
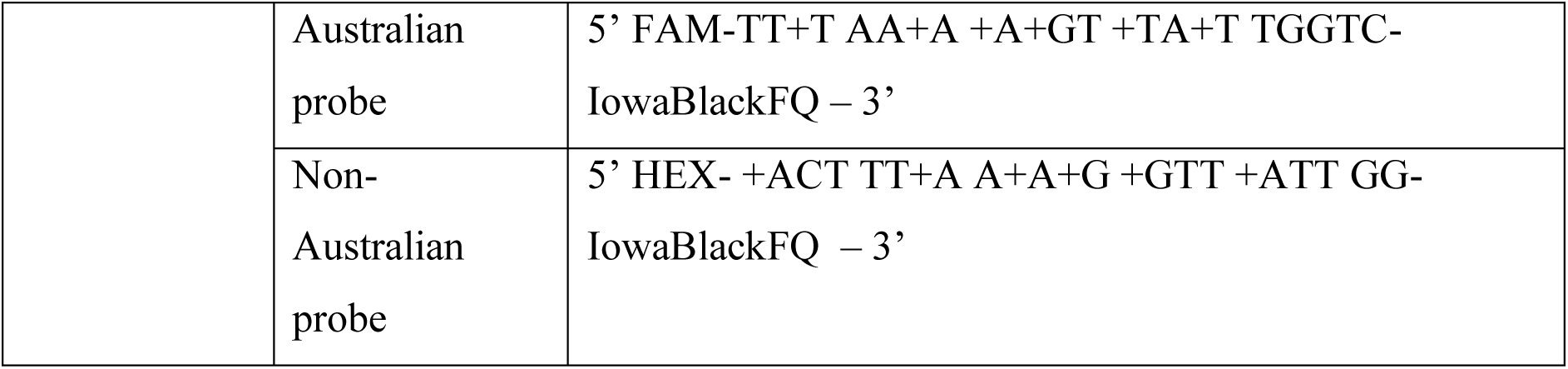
Primers used in this study. Locked Nucleic Acid (LNA) bases are denoted by a + symbol on the left side of the base.

**Supplementary Table 2.** Validation of CO1 and CO2 mitotyping assays against wild-type populations.

| Primer set | Wild-type population | No. mosquitoes with non-founder SNP | No. mosquitoes with founder SNP | % wild-type population with founder SNP |
| --- | --- | --- | --- | --- |
| **CO1** | Vanuatu | 401 | 3 | 0.75% |
|  | Fiji | 476 | 0 | 0% |
|  | Kiribati | 365 | 1 | 0.28% |
|  | Colombia | 41 | 0 | 0% |
| **CO2** | Brazil | 40 | 0 | 0% |

**Supplementary Table 3:**
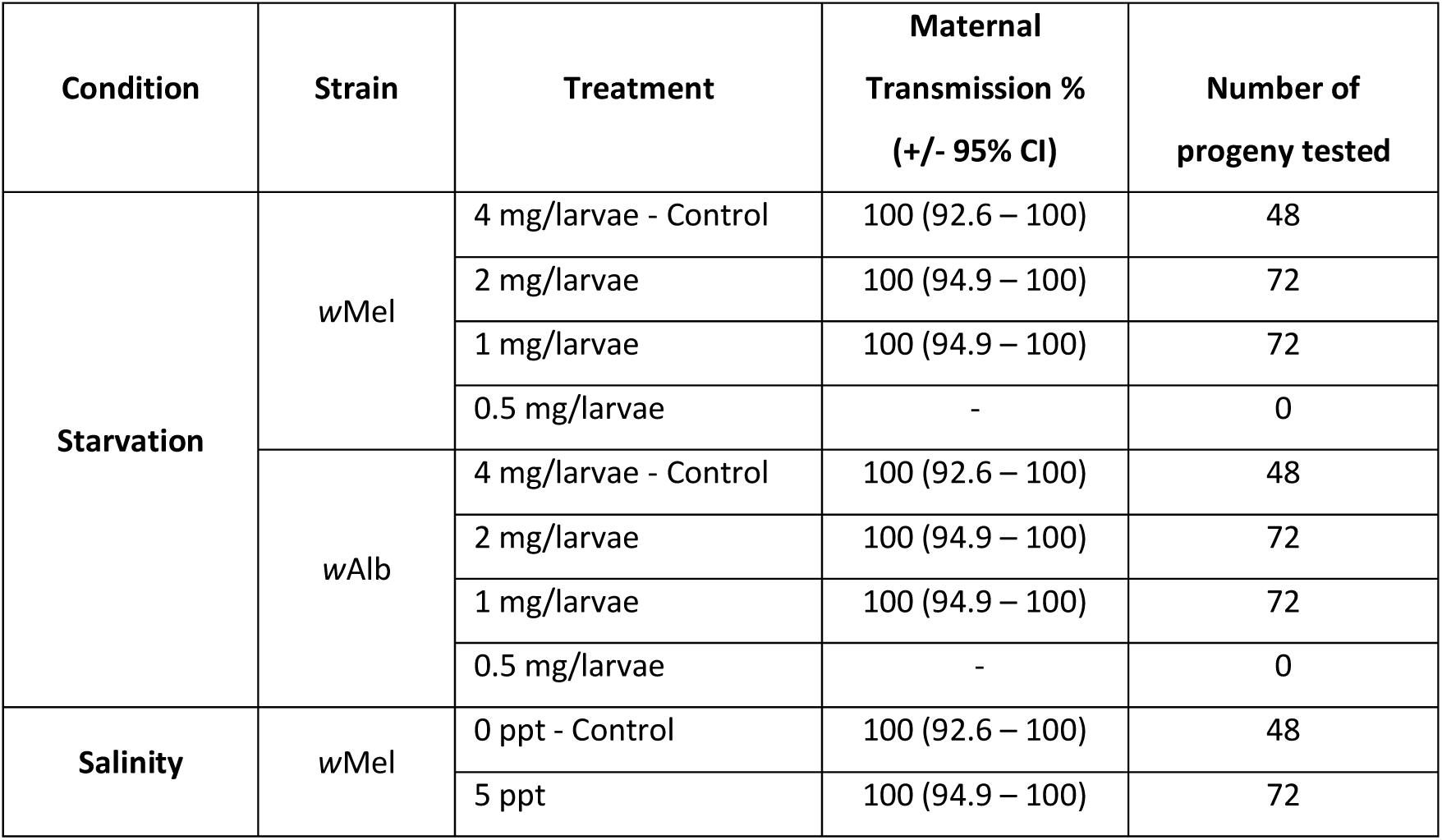
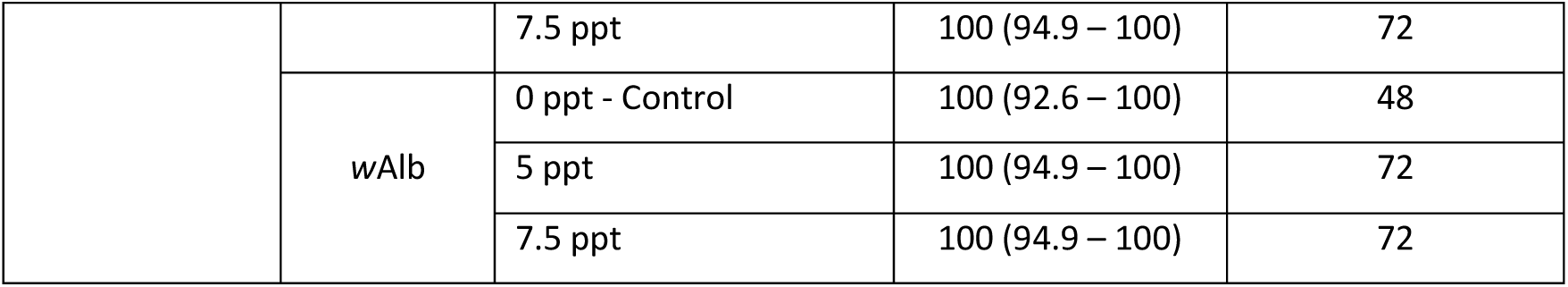
Maternal transmission of *Wolbachia* in *Ae. aegypti* post-exposure. DPE – days post emergence, ppt – parts per thousand.

